# PD-L1 blockade restores CAR T cell activity through IFNγ-regulation of CD163+ macrophages

**DOI:** 10.1101/2022.01.25.477150

**Authors:** Yukiko Yamaguchi, Jackson Gibson, Kevin Ou, Rachel H. Ng, Neena Leggett, Vanessa D. Jonsson, Jelani C. Zarif, Peter P. Lee, Xiuli Wang, Catalina Martinez, Tanya B. Dorff, Stephen J. Forman, Saul J. Priceman

**Author notes:** To whom correspondence may be addressed: Saul J. Priceman, Department of Hematology and Hematopoietic Cell Transplantation, City of Hope, 1500 E. Duarte Rd, Duarte CA 91010, USA.

## Abstract

**Background:** The immune suppressive tumor microenvironment (TME) that inhibits T cell infiltration, survival, and anti-tumor activity has posed a major challenge for developing effective immunotherapies for solid tumors. Chimeric antigen receptor (CAR)-engineered T cell therapy has shown unprecedented clinical response in treating patients with hematological malignancies, and intense investigation is underway to achieve similar responses with solid tumors. Immunologically cold tumors, including prostate cancers, are often infiltrated with abundant tumor-associated macrophages (TAMs), and infiltration of CD163^+^ M2 macrophages correlates with tumor progression and poor responses to immunotherapy. However, the impact of TAMs on CAR T cell activity alone and in combination with TME immunomodulators is unclear.

**Methods:** To model this *in vitro*, we utilized a novel co-culture system with tumor cells, CAR T cells, and polarized M1 or M2 macrophages from CD14^+^ PBMCs collected from healthy human donors. Tumor cell killing, T cell activation and proliferation, and macrophage phenotypes were evaluated by flow cytometry, cytokine production, RNA sequencing, and functional blockade of signaling pathways using antibodies and small molecule inhibitors. We also evaluated the TME in humanized mice following CAR T cell therapy for validation of our *in vitro* findings.

**Results:** We observed inhibition of CAR T cell activity with the presence of M2 macrophages, but not M1 macrophages, coinciding with a robust induction of PD-L1 in M2 macrophages. We observed similar PD-L1 expression in TAMs following CAR T cell therapy in the TME of humanized mice. PD-L1, but not PD-1, blockade in combination with CAR T cell therapy altered phenotypes to more M1-like subsets and led to loss of CD163^+^ M2 macrophages via IFNγ signaling, resulting in improved anti-tumor activity of CAR T cells.

**Conclusion:** This study reveals an alternative mechanism by which the combination of CAR T cells and immune checkpoint blockade modulates the immune landscape of solid tumors to enhance therapeutic efficacy of CAR T cells.

## Introduction

Adoptive transfer of chimeric antigen receptor (CAR)-engineered T cells has demonstrated robust and durable clinical efficacy in patients with B-cell malignancies,[1-3], but to date has shown underwhelming response rates in patients with solid tumors,[4, 5]. This clinical observation is in large part attributed to the immune-suppressive tumor microenvironment (TME) of solid tumors, comprising infiltrating myeloid cells and regulatory T cells that inhibit endogenous anti-tumor immunity and adoptively transferred cell therapies. Overcoming this challenge will be critical to unleashing the full potential for CAR T cell therapies for solid tumors, and likely will require disease- and context-specific considerations.

Tumor-associated macrophages (TAMs) are the most abundant immune cells in many solid tumors, and TAM infiltration strongly correlates with tumor progression and poor prognosis in various solid tumors,[6-10] and lymphoma,[11]. While macrophages retain phenotypic and functional plasticity, the majority of TAMs are immune-suppressive, M2-like macrophages with complex pro-tumor functions. TAMs secrete various cytokines and growth factors including IL-10, TGFβ, VEGF, and CXCL12 to drive cancer progression through immune suppression, tumor angiogenesis, invasion and metastasis,[12-14]. TAMs also play critical roles in response and resistance to common cancer therapies such as chemotherapy, radiation therapy,[15], angiogenesis,[16] and hormone deprivation therapy,[17], and numerous macrophage-modulating approaches have shown improved therapeutic efficacy in preclinical studies,[12, 18-21].

Preclinical studies also demonstrated that TAMs mediate resistance to immune checkpoint blockade (ICB),[22-24], and targeting TAMs likely alter outcomes of clinical interventions,[25]. PD-1 and PD-L1 are expressed in various immune cells including T cells,[26], NK cells,[27] and macrophages,[28]. Tumor PD-L1 expression does not accurately predict clinical response to anti-PD-L1 therapy, and more recent studies indicate that PD-L1 expressed by immune cells may contribute to immune suppression,[27-29]. Macrophage PD-L1 is particularly abundant in the TME, but the role of PD-L1 signaling in macrophages and the direct impact of anti-PD-L1 blockade on macrophages remains controversial,[28-30]. Recent studies have shown that CAR T cells, especially in combination with other therapeutic agents, modulate myeloid cell phenotypes and alter the immune-suppressive TME,[31-33]. ICB has been utilized in combination with CAR T cell therapy, with the notion that induction of immune responses with CAR T cells may instigate checkpoint pathways in immunologically cold tumors as a compensatory resistance mechanism, providing rationale for the therapeutic combination. Despite a clinical need for overcoming immune suppression to improve CAR T cell therapies for solid tumors, preclinical modeling of this phenomenon is complicated and remain limited in their predictive capabilities.

In this study, we aimed to develop an *in vitro* model to recapitulate the suppression of CAR T cells in microenvironments with abundant immune-suppressive macrophages. In this model system, target tumor cells and CAR T cells were co-cultured in the presence of M1- or M2-polarized macrophages to evaluate their respective roles in CAR T cell functionality. We showed that M1 macrophages promote, while M2 macrophages suppress, CAR T cell-mediated tumor cell killing and cytokine production. We also observed CAR T cell-regulated PD-L1 induction in both tumor cells and macrophages *in vitro*, with induction levels found to be most dramatic in M2 macrophages. We confirmed CAR T cell-regulated PD-L1 induction in TAMs using an *in vivo* humanized mouse model of prostate cancer. By blocking PD-L1 with atezolizumab or avelumab, we found that inhibiting macrophage PD-L1 was sufficient to restore CAR T cell-mediated tumor killing. However, this restoration of CAR T cell killing by blockade of PD-L1 appears independent of canonical PD-1/PD-L1 signaling, as the phenomenon was not seen with blockade of PD-1 with nivolumab. Instead, PD-L1 inhibition specifically and potently depleted M2 macrophages in the presence of CAR T cells. These findings give mechanistic insights by which CAR T cell and ICB combination therapies enhance anti-tumor immunity in an immune-suppressive TME and is a useful model to study macrophage-mediated immune suppression.

## Results

### Human Monocyte-Derived M2 Macrophages Suppress CAR T Cells *In Vitro*

Macrophages are an abundant immune cell population in lymphoma,[11] and various solid tumors including prostate cancer,[7, 8], and their abundance correlates with metastasis and poor prognosis. To investigate the impact of macrophage-rich immunosuppressive solid tumor microenvironment (TME) on CAR T cells, we developed an *in vitro* immune-suppression assay by co-culturing CAR T cells, M1 or M2 macrophages and target tumor cells at an effector:macrophage:tumor ratio of 1:5:10 (**Figure 1a**). Macrophages were differentiated from CD14^+^ cells enriched from healthy donor PBMCs and *in vitro* polarized as previously described,[34] into M1 (CD80^high^, CD163^-^, CD206^low^) or M2 (CD80^low^, CD163^+^, CD206^high^) macrophages (**Figure S1a, b**). To model the prostate TME, DU145 prostate tumor cells were engineered to express prostate stem cell antigen (PSCA) and co-cultured with untransduced (UTD) or PSCA-CAR T cells previously developed by our group,[35]. CD19-CAR T cells,[36] and Daudi lymphoma cells were used to model the lymphoma TME. We evaluated antitumor activity, activation and proliferation of CAR T cells by flow cytometry (**Figure S2**) and interferon-γ (IFNγ) secretion by ELISA. CAR T cell anti-tumor activity was normalized to UTD T cells, and activation was measured by 4-1BB upregulation. In both prostate and lymphoma models, anti-tumor cytolytic activity of T cells was inhibited in the presence of M2 macrophages, while it was enhanced in the presence of M1 macrophages (**Figure 1b-d**). T cell proliferation (**Figure 1e**), activation (**Figure 1f, g, Figure S3**), and IFNγ secretion (**Figure 1h**) were also inhibited by M2 macrophages. Collectively, these data show that our *in vitro* co-culture system effectively recapitulates the immunosuppressive effects of M2 macrophages on CAR T cells in the TME.

**Figure 1:**
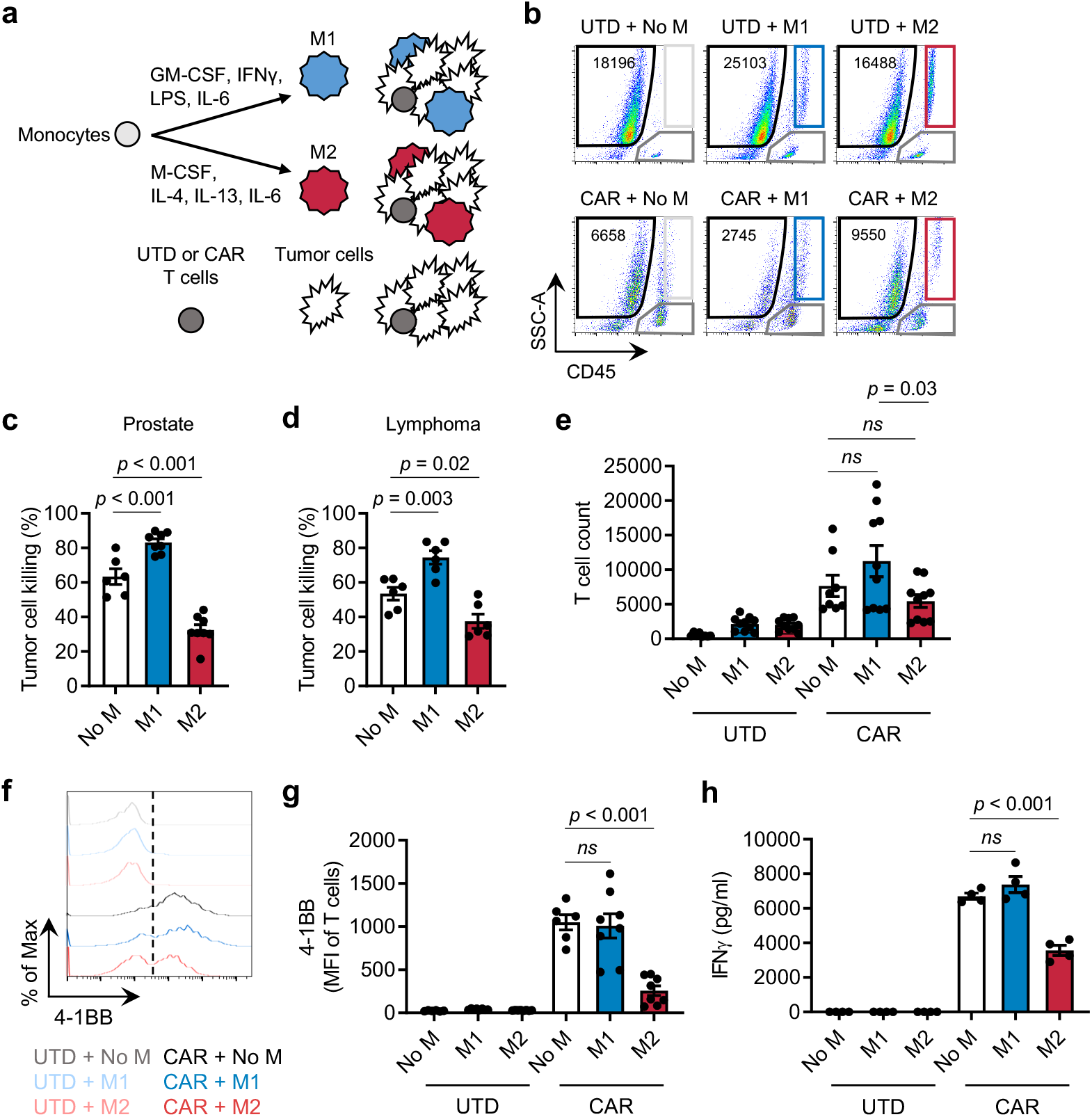
M2 macrophages suppress CAR T cells. (a) Illustration of the immune-suppression assay. CD14^+^ PBMCs were differentiated and polarized to M1 or M2 macrophages *in vitro*, and macrophages, CAR T cells, and tumor cells were co-cultured and evaluated for functional activities by flow cytometry. (b) Flow cytometry plots indicating the number of viable tumor cells in each condition. (c, d) CAR T cell-mediated tumor cell killing of DU145-PSCA prostate cancer (c) and CD19^+^ Daudi lymphoma (d) cells in the presence or absence of M1 or M2 macrophages after 6 and 3 days, respectively. CAR T cell-mediated tumor cell killing was normalized to untransduced (UTD) T cells. (e-h) Proliferation (10 days) (e), 4-1BB activation (6 days) (f, g), and IFNγ secretion (3 days) (h) of T cells in the presence or absence of M1 or M2 macrophages in the prostate cancer model. Proliferation and activation of T cells was measured by flow cytometry. Secreted IFNγ in supernatant was measured by ELISA.

### CAR T Cells Alter the Phenotype of M2 Macrophages *In Vitro*

Next, we investigated the impact of CAR T cells on the TME by evaluating phenotypic changes that CAR T cells induce in macrophages. In the immune-suppression assay, we assessed expression of CD80 and CD163 as classical M1 and M2 markers in M2 macrophages in the presence or absence of PSCA-CAR T cells by flow cytometry (**Figure 2a**). We found that CAR T cells upregulated CD80 (**Figure 2b**) and downregulated CD163 **(Figure 2c**) surface expression on M2 macrophages. To evaluate whether such phenotypic changes are mediated by secreted factors, we collected conditioned media from tumor killing assay where DU145-PSCA tumor cells were co-cultured with PSCA-CAR T cells (**Figure 2d**). The conditioned media was applied onto adherent M2 macrophages, and their phenotype was assessed after 48 hours. Phenotypic changes induced in M2 macrophages mirrored the observation in the immune-suppression assay (**Figure 2e, f**), suggesting that CAR T cells alter M2 macrophage phenotype via secreted factors. Furthermore, transcriptome analysis of M2 macrophages by bulk RNA-seq revealed a global gene expression change upon stimulating with the CAR T cell-derived CM, and M1 signatures including CD80, CXCL9 and IL1B increased while M2 signatures including CD163, ADORA3 and IL10 decreased (**Figure 2g**). We found by gene ontology analysis that inflammatory pathways were activated (**Figure 2h**), further supporting changes that CAR T cells induce in M2 macrophages. CAR T cell-derived conditioned media did not induce phenotypic changes in M1 macrophages (**Figure S4**). Taken together, these results indicate that CAR T cells alter the TME by repolarizing M2 macrophages to a less immune suppressive, M1-like macrophage state via paracrine signaling.

**Figure 2:**
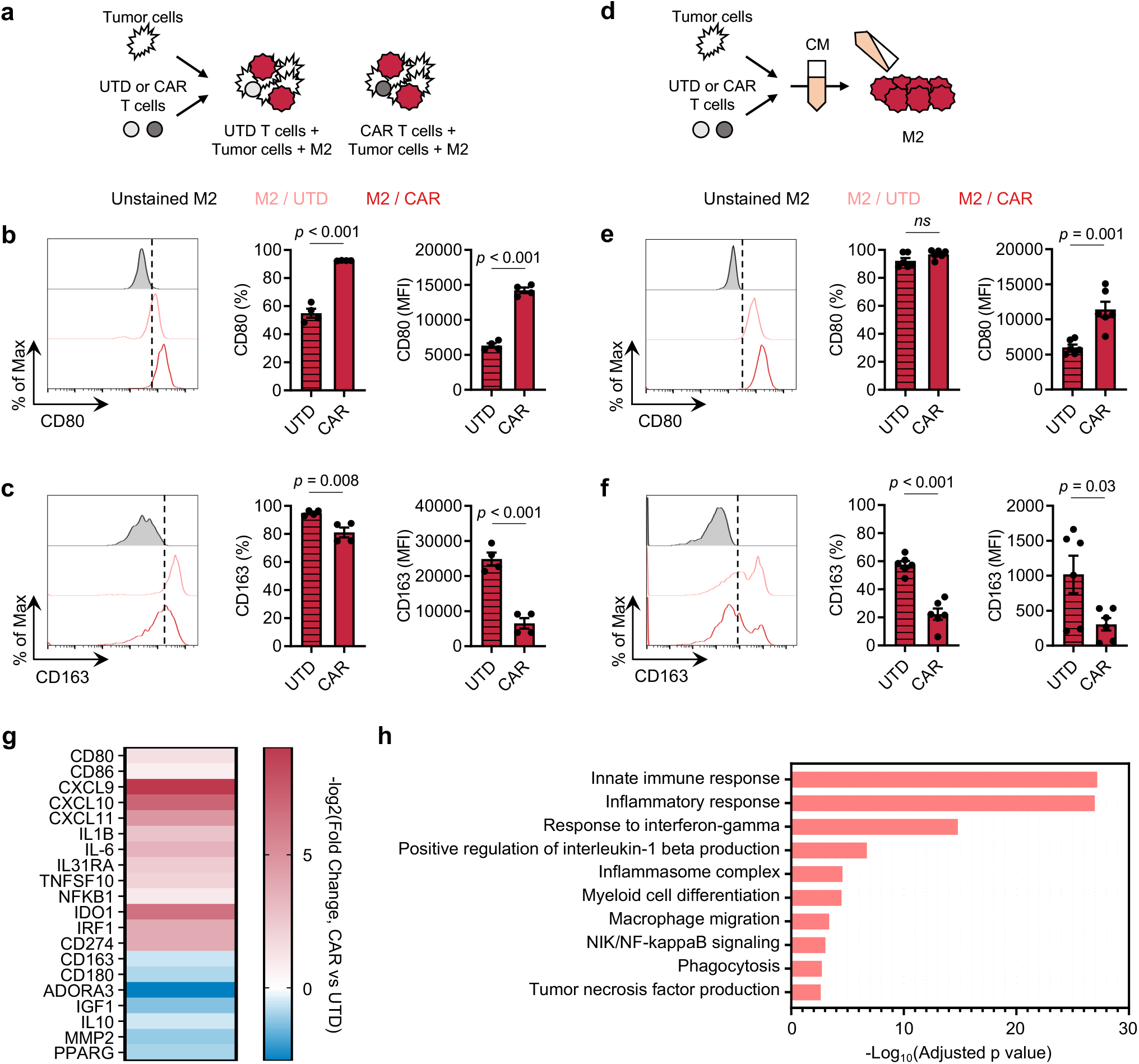
CAR T cells alter M2 macrophage phenotypes. (a) Illustration of the immune-suppression assay to evaluate M2 macrophage phenotype. (b, c) Cell surface expression of CD80 (b) and CD163 (c) in M2 macrophages in the immune-suppression assay evaluated by flow cytometry. (d) Illustration of M2 macrophage stimulation with conditioned media (CM) derived from CAR T cell:tumor cell co-cultures. (e, f) Cell surface expression of CD80 (e) and CD163 (f) in M2 macrophages evaluated by flow cytometry 48 hours after CM stimulation. (g) Transcriptional changes by bulk RNA-seq induced in M2 macrophages upon stimulation with CAR T cell-derived CM. Expression of selected immune-related genes is shown relative to a control condition stimulated with UTD T cell-derived CM. (h) Gene ontology (GO) enrichment analysis highlighting activated immune-related biological pathways in M2 macrophages upon stimulation with CAR T cell-derived CM.

### PD-L1 is upregulated in M2 macrophages in the presence of CAR T cells

IFNγ is a well-known inducer of programmed death-ligand 1 (PD-L1) and one of the cytokines T cells secrete upon activation and has been suggested to be a pathway of resistance to cellular immunotherapy,[28, 37]. Therefore, we assessed PD-L1 expression changes in M2 macrophages and tumor cells in the immune-suppression assay. In the prostate model, both DU145-PSCA tumor cells and M2 macrophages induced PD-L1 surface expression in the presence of CAR T cells. Interestingly, M2 macrophages showed greater induction in frequency and abundance of PD-L1 expression compared to tumor cells and M1 macrophages (**Figure 3a-c**). In the lymphoma model, PD-L1 was induced in M2 macrophages (**Figure S5a**) but not in Daudi tumor cells (**Figure S5b**). We hypothesized that PD-L1 was induced in a paracrine fashion, and to test this hypothesis, we treated M2 macrophages and various tumor cells with conditioned media obtained from tumor killing assays. PD-L1 induction was the greatest in M2 macrophages at the protein (**Figure S6a, b**) and mRNA levels (**Figure S6c**), recapitulating induction in the *in vitro* immune suppression assay.

**Figure 3:**
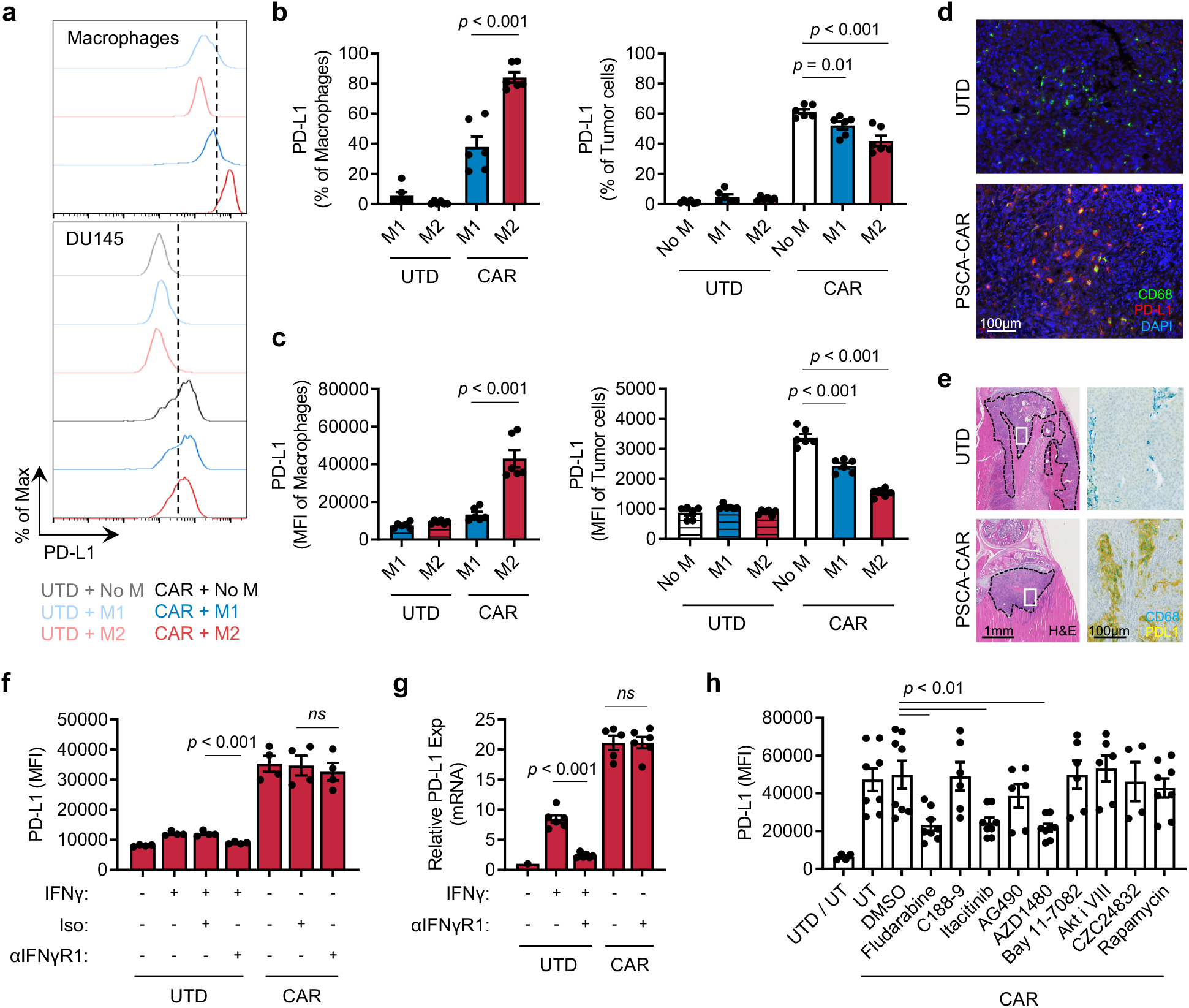
CAR T cells induce PD-L1 expression in M2 macrophages. (a-c) PD-L1 expression in macrophages and DU145-PSCA tumor cells in the immune-suppression assay. (d, e) Immunostaining of CD68 and PD-L1 in a humanized MISTRG mouse model following CAR T cell therapy against subcutaneous DU145-PSCA (d) and intratibial LAPC9 (e) prostate xenografts. (f, g) PD-L1 induction at the protein (f) and mRNA (g) levels following inhibition of IFNγ signaling. Anti-IFNγR1 antibody was used to block IFNγ signaling in the presence of recombinant IFNγ or CAR T cell-derived CM collected from the tumor cell killing assay. (h) PD-L1 induction following inhibition of various signaling pathways. CAR T cell-derived CM was applied to M2 macrophages in the presence of various small molecule inhibitors: fludarabine (STAT1 i), C188-9 (STAT3 i), itacitinib (JAK1 i), AG490 (JAK2 i), AZD1480 (JAK1/2 i), Bay11-7082 (NFκB i), Akti VIII (AKT i), CZC24832 (PI3K i), rapamycin (mTOR i). PD-L1 induction was evaluated by flow cytometry 48 hours after CM stimulation.

To evaluate whether CAR T cells induce PD-L1 expression in tumor-associated macrophages *in vivo*, we humanized mice by engrafting human CD34^+^ hematopoietic stem cells in immune-compromised MISTRG mice,[38]. DU145-PSCA tumor cells were then injected subcutaneously, and LAPC9 cells, a patient-derived metastatic prostate cancer cell line with endogenous PSCA expression, were injected into the intratibial space to model bone metastatic disease,[35]. PSCA-CAR T cells were adoptively transferred via intravenous injection, as we have done previously in our preclinical therapeutic studies,[35]. In humanized MISTRG mice, CD68^+^ human macrophages efficiently infiltrated into human tumor xenografts, and immunostaining revealed colocalization of CD68 and PD-L1 in DU145-PSCA (**Figure 3d**) and in LAPC9 (**Figure 3e**) xenograft. These data, collectively, show that CAR T cells directly induce PD-L1 in both tumor cells and M2 macrophages *in vitro* and *in vivo*.

### IFNγ is not a dominant inducer of PD-L1 expression by CAR T cells

We next hypothesized that PD-L1 is induced by IFNγ, and to test the hypothesis, we treated M2 macrophages and DU145 tumor cells with conditioned media collected from the tumor killing assay in the presence of anti-IFNγR1 antibody. Cells were collected after 48 hours to evaluate PD-L1 protein expression by flow cytometry (**Figure 3f, Figure S7a**), and cell lysates were collected after 6 hours to measure mRNA expression by qPCR (**Figure 3g, Figure S7b**). Blocking IFNγ signaling was not sufficient to inhibit PD-L1 expression in M2 macrophages or DU145 tumor cells in the conditioned media. Also, recombinant IFNγ only moderately induced PD-L1 expression when it was added at similar concentrations (~20 ng/ml) measured in CAR T cell-derived conditioned media (**Figure S8a**). Increasing the concentration of recombinant IFNγ up to 200 ng/ml did not reach the level of PD-L1 induction in M1 and M2 macrophages observed with CAR T cell-conditioned media (**Figure S8b, c**). We treated M1 and M2 macrophages with varying concentrations of conditioned media and showed that 5-20% conditioned media was sufficient to induce maximal levels of PD-L1 (**Figure S8d**). Despite IFNγ being a well-established PD-L1 inducer, these results indicate that IFNγ is not a sole or dominant inducer of PD-L1 expression in tumor cells or M2 macrophages in this system. The data suggest that PD-L1 induction is regulated by the presence of other inducers in CAR T cell-derived soluble factors.

To identify signaling pathways that mediate PD-L1 induction, we treated M2 macrophages with small molecule inhibitors of various pathways. While inhibition of STAT3, NFκB, AKT, PI3K and mTOR signaling was not sufficient to block PD-L1 induction by CAR T cells in M2 macrophages, inhibition of STAT1 with fludarabine resulted in loss of PD-L1 induction in M2 macrophages (**Figure 3h**). Loss of PD-L1 induction was also shown following JAK1/2 inhibition with AZD1480 as well as JAK1-selective inhibition with itacitinib, but not by JAK2 inhibition with AG490. These results indicate that PD-L1 expression induced by CAR T cells is mediated primarily by a JAK1/STAT1 pathway, independent of IFNγ.

### PD-L1 blockade inhibits M2 macrophage-mediated suppression of CAR T cells

To test the functionality of PD-L1 in immune suppression by M2 macrophages, we blocked PD-L1 with atezolizumab, an anti-PD-L1 monoclonal antibody, in the *in vitro* immune suppression assay. PD-L1 blockade restored CAR T cell-mediated tumor cell killing in the presence of M2 macrophages (**Figure 4a, b**). This observation was reproduced using avelumab, another anti-PD-L1 monoclonal antibody (**Figure 4e**). T cell activation (**Figure 4c**) and IFNγ secretion (**Figure 4d**) were also restored in the presence of PD-L1 blockade, supporting a role for PD-L1 in regulating M2 macrophage-mediated immune suppression. Furthermore, tumor killing was also restored in the lymphoma model (**Figure S9**) where Daudi tumor cells lacked PD-L1 expression (**Figure S5b**). These data indicate that macrophage PD-L1 is sufficient to drive immune suppression. However, blocking PD-1 using Nivolumab, an anti-PD-1 monoclonal antibody, did not restore CAR T cell-mediated tumor cell killing in a similar fashion (**Figure S10a**). Interestingly, consistent with our previous publication,[35], PD-1 was not readily induced in CAR T cells (**Figure S10b**). Therefore, while M2 macrophage PD-L1 is necessary for immune suppression in this system, these results indicated that the classical PD-1/PD-L1 signaling axis is not a primary mechanism by which M2 macrophages suppress CAR T cells.

**Figure 4:**
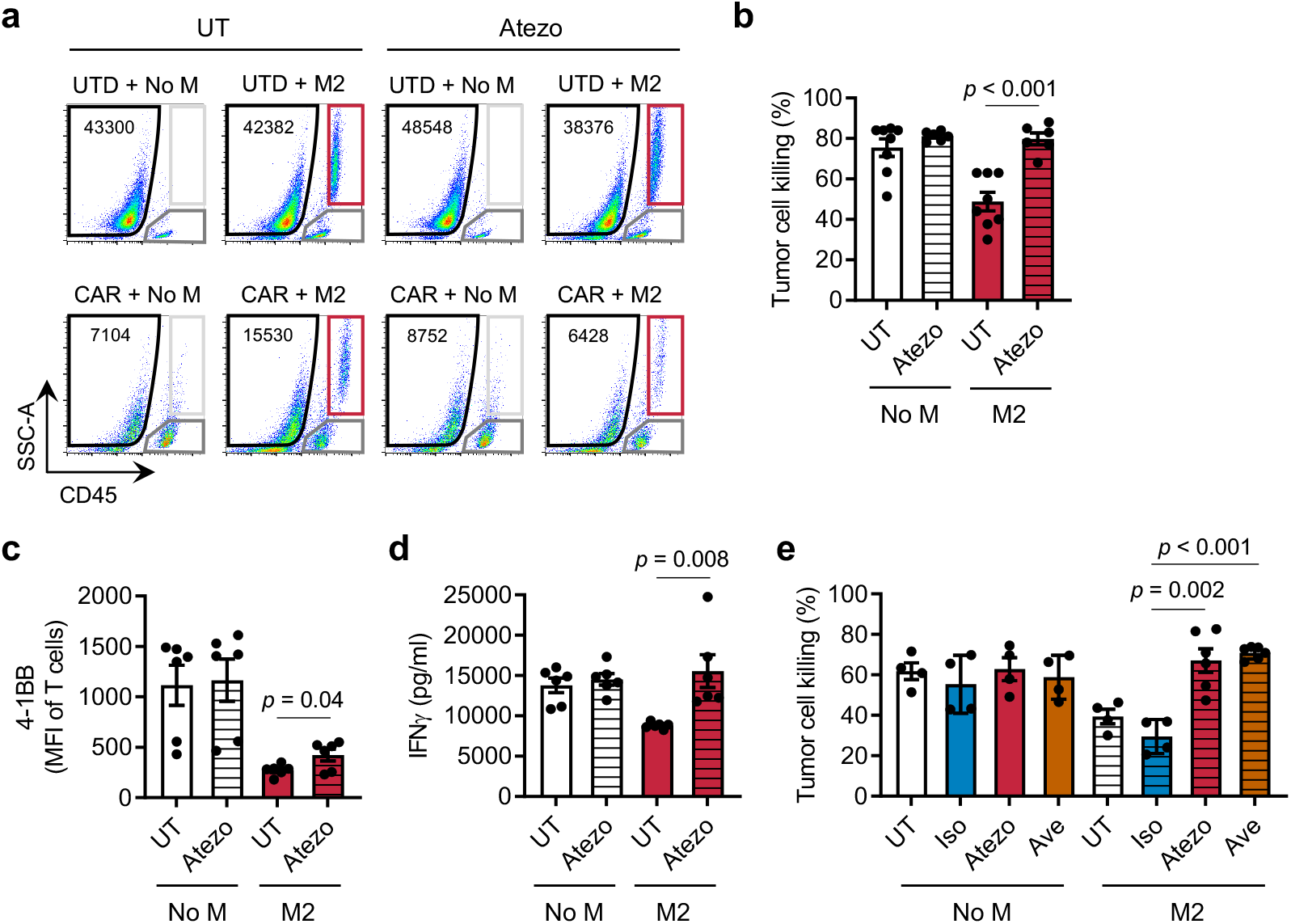
PD-L1 blockade restores CAR T cell function in the presence of suppressive M2 macrophages. CAR T cell function was evaluated in the immune-suppression assay in the presence of PD-L1 blockade. (a) Flow cytometry plots indicating the number of viable tumor cells in each condition in the presence or absence of anti-PD-L1 antibody, Atezolizumab (Atezo). (b) Quantification of CAR T cell-mediated killing of DU145-PSCA tumor cells in the presence or absence of Atezo. (c) 4-1BB T cell activation was evaluated by flow cytometry. (d) IFNγ secretion was measured by ELISA. (e) Tumor cell killing of CAR T cells in the presence or absence of two clinically approved anti-PD-L1 antibodies, Atezo and Avelumab (Ave).

### Combining CAR T cells and PD-L1 blockade alter phenotype and reduce survival of M2 macrophages

Macrophages express PD-1 and PD-L1 (**Figure S1b)**, and increasing evidence supports that these cell surface receptors play a role in shaping intrinsic cellular properties of macrophages including their immune suppressive function,[30, 39, 40]. We hypothesized that blocking PD-L1 alters the ability of M2 macrophages to suppress CAR T cells. First, we assessed M2 macrophages in the immune-suppression assay in the presence of PD-L1 blockade. In the presence of CAR T cells, the number of M2 macrophages decreased compared to respective controls with UTD T cells (**Figure 4a, Figure 5a**). The combination of CAR T cells and PD-L1 blockade resulted in significantly fewer M2 macrophages, and specifically in the presence of CAR T cells. These data suggest that PD-L1 blockade specifically in combination with CAR T cells has a direct impact on survival of M2 macrophages. We also evaluated macrophage phenotype and found fewer CD163^+^ M2 macrophages in the combination of CAR T cells and PD-L1 blockade (**Figure 5b**). To further interrogate the mechanism underlying this phenomenon, we stimulated M2 macrophages with conditioned media collected from tumor-CAR T cell co-cultures. Consistent with the previous observation in the immune suppression assay, the combination of CAR T cells and PD-L1 blockade resulted in reduction of total viable and CD163+ M2 macrophages (**Figure 5c, d**). Given previous studies in the field suggesting the importance of PD-1 in immune suppression by macrophages,[39, 40], we also blocked PD-1 with Nivolumab, but did not observe a similar impact on viability or changes in CD163 expression in M2 macrophage (**Figure 5c, d**).

**Figure 5:**
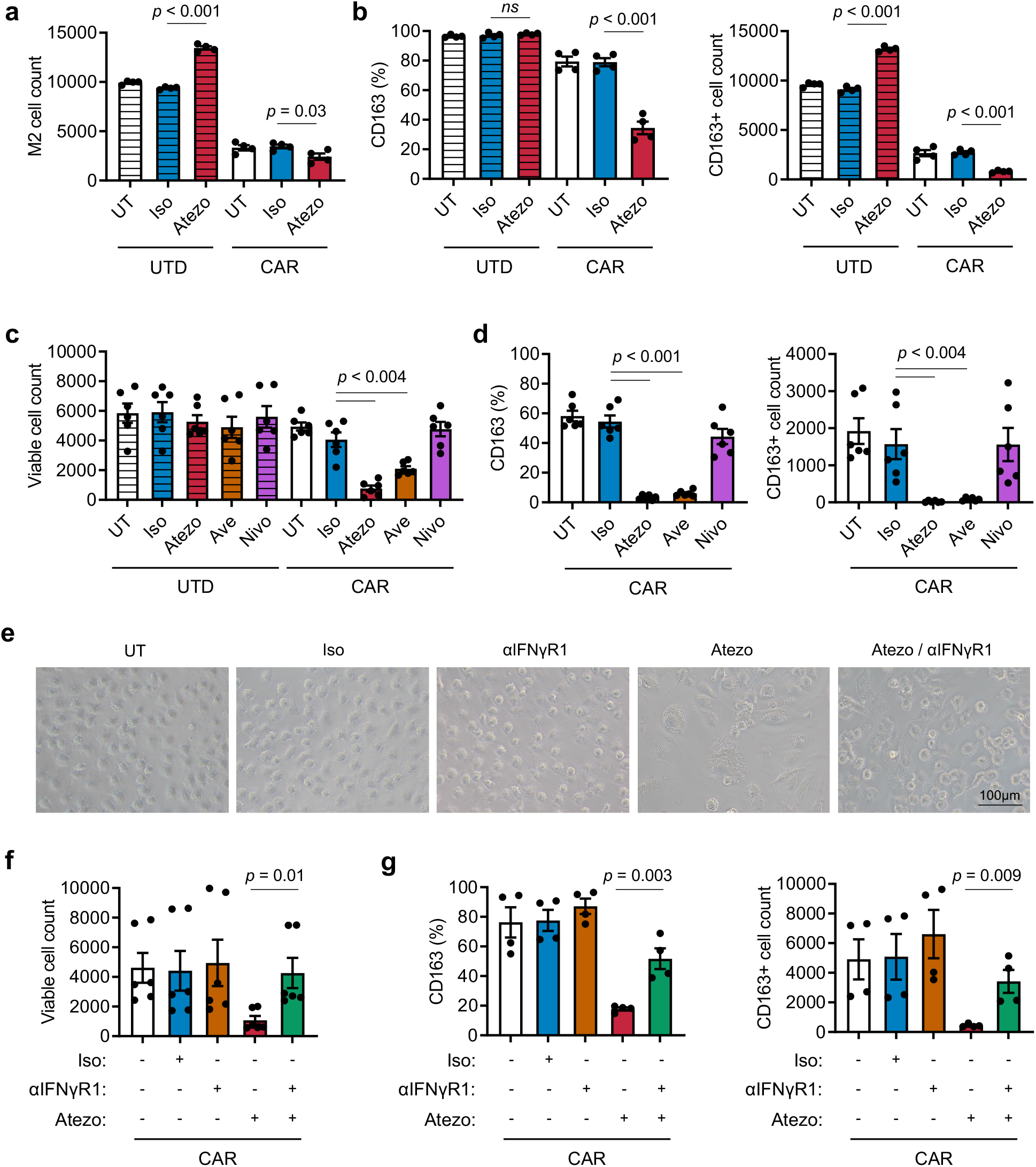
Combination of PD-L1 blockade and CAR T cell therapy depletes M2 macrophages via IFNγ signaling. (a, b) Analysis of M2 macrophages in the immune-suppression assay in the presence or absence of PD-L1 blockade. (c, d) Analysis of M2 macrophages stimulated with CAR T cell-derived CM in the presence or absence of PD-1 or PD-L1 blockade. (e-g) Images and analysis of M2 macrophage stimulated with CAR T cell-derived CM in the presence or absence of PD-L1 and/or IFNγR1 blockade. (f, g) The number of total viable M2 macrophages (a, c, f) and the frequency and number of CD163^+^ M2 macrophages (b, d, g) were evaluated by flow cytometry.

### IFNγ signaling mediates altered phenotype of M2 macrophages following PD-L1 inhibition

FNγ activates macrophages and plays important roles in promoting inflammation. Therefore, we hypothesized that IFNγ mediates the loss of CD163^+^ M2 macrophages in the combination of CAR T cells and PD-L1 blockade. To test this, we treated M2 macrophages with anti-PD-L1 and anti-IFNγR1 antibodies in the conditioned media collected from tumor-CAR T cell co-cultures. By microscopy, we not only visually confirmed the reduction in M2 macrophage cell numbers with PD-L1 inhibition, but also observed M2 macrophages become enlarged and vacuolated (**Figure 5e**). Blocking IFNγ signaling prevented these morphological changes and loss of CD163^+^ cells induced by PD-L1 inhibition (**Figure 5e, f, g**), suggesting that the combination of CAR T cells and PD-L1 blockade directly impacts M2 macrophages via IFNγ signaling, reversing M2 macrophage-mediated immunosuppression of CAR T cells.

## Discussion

In the current study, we investigated the impact of myeloid cells on CAR T cell activity using an *in vitro* model that we established to recapitulate the immune-suppressive TME. We found that M2 macrophages, but not M1 macrophages, suppressed the anti-tumor activity of CAR T cells using both PSCA^+^ prostate cancer and CD19^+^ lymphoma models. The presence of CAR T cells altered the phenotype of M2 macrophages towards a less immune-suppressive state with reduced M2-like CD163^+^ and greater M1-like CD80^+^ populations. We also observed induction of PD-L1 expression in tumor cells as well as M1 and M2 macrophages, but M2 macrophages had significantly higher cell-surface density of PD-L1 induction than in tumor cells or M1 macrophages. Inhibition of PD-L1 using antibody blockade restored CAR T cell function suppressed by M2 macrophages, but this restoration was not mediated by canonical PD-1/PD-L1 axis as CAR T cell function was not restored with PD-1 blockade. Instead, the combination of CAR T cells and PD-L1 blockade resulted in fewer CD163^+^ M2 macrophages, suggesting a direct impact on these cells. Further, we showed that IFNγ was required for this phenomenon, as inhibition of IFNγR signaling potently reversed this PD-L1-regulated survival of M2 macrophage. These findings provide mechanistic insight into CAR T cell-mediated alterations in the TME and specifically on immune-suppressive myeloid cells. However, our studies suggest CAR T cells alone may not be sufficient to overcome immunosuppression in the TME and may require PD-L1 blockade to enable the full therapeutic potential of CAR T cells.

While recent evidence supports the notion that CAR T cells alone can enhance endogenous immunity, numerous studies have shown that CAR T cell therapy is not able to elicit adequate clinical response against solid tumors,[41, 42], justifying rational for combining immunotherapies. Our *in vitro* model confirms the ability of CAR T cells to alter the myeloid cell subsets to a less suppressive state, but such immunomodulation was not sufficient for CAR T cells to evade immune suppression. Moreover, we observed this M2 macrophage shift to a more pro-inflammatory state in approximately 60% of tested healthy human donors, demonstrating apparent heterogeneity in CAR T cell-mediated immunomodulation and susceptibility of macrophages among individuals. Studies in mouse models might reproduce immunomodulation of macrophages in response to CAR T cells, but the use of inbred mice might not adequately uncover heterogenous responses that we observed in our *in vitro* model. We may be able to use this model in the future to better understand and develop therapies that enhance how CAR T cells function in the presence of TMEs with abundant M2 macrophage subsets as seen in prostate cancers and other solid tumors.

PD-1/PD-L1 blockade combined with CAR T cells is a current clinical strategy owing largely to the field’s collective evidence that immune checkpoint pathways are induced following activity of CAR T cells, which may ultimately lead to exhaustion of CAR T cells. The contribution of myeloid PD-L1 expression on immunosuppression within the tumor microenvironment has been reported in preclinical models and could be regulated by alternative mechanisms,[28, 30]. In our study, the canonical PD-1/PD-L1 axis did not directly influence CAR T cell function, as PD-L1 blockade, but not PD-1 blockade, reversed macrophage-mediated immune suppression. Our data suggest that CAR T cell-mediated PD-L1 expression in macrophages may specifically and directly drive their survival and immune-suppressive phenotype. The change in CD163 expression of macrophages in response to CAR T cells was variable among individuals, however, combining CAR T cells with PD-L1 blockade induced a uniform response in all tested individuals. Loss of immune-suppressive macrophages with the combination of CAR T cells and PD-L1 blockade resembles observations with other myeloid-targeting therapies, including CSF1/CSF1R blockade^14,36^, CCL2/CCR2 inhibition,[13, 20] and novel anti-CD206 peptides,[18]. Due to this mechanism of action of TME remodeling, the efficacy of combining CAR T cells and PD-L1 blockade may be driven in part by tumor composition and density of macrophages. Our data suggest that this combination therapy may be more effective in immunologically “cold” solid tumors with abundant CD163^+^ immune suppressive macrophages.

The requirement of IFNγ in regulating the survival and function of M2 macrophage following PD-L1 blockade suggests that amplifying IFNγ signaling may be an actionable target for improving the combination of CAR T cells and ICB. Various engineering and manufacturing approaches can enhance IFNγ secretion by CAR T cells,[35, 36, 43, 44], and CAR T cells with greater IFNγ secretion may better remodel the TME in combination with ICB. Although we found that IFNγ was critical for PD-L1 blockade-induced M2 macrophage depletion, mechanisms of how the combination impact functions of immune suppressive macrophages remains unclear. Although increased apoptosis of CD163^+^ cells in the combination of CAR T cells and PD-L1 blockade was expected, we failed to demonstrate increased apoptosis in our studies. Using time-lapsed imaging, we revealed cells pursuing and catching adjacent cells before morphological changes occurred, indicating possible antibody-dependent cellular phagocytosis of M2 macrophages. Also, macrophages are known to enlarge and form vacuoles via fusion in chronic inflammation,[45], and the morphological changes may be a manifestation of a highly inflammatory state. Further studies are warranted to elucidate mechanisms of immunomodulation that macrophages undergo following CAR T cell therapy and PD-L1 blockade.

We built the immune suppression assay under an assumption that TAMs are M2-like, immune suppressive macrophages. However, macrophages phenotypes and functions are not as binary as M1 or M2, but rather demonstrate plasticity along a spectrum of phenotypes and functions. In addition to macrophage cell plasticity, the disease context and clinical interventions likely contribute to shaping the phenotype of TAMs. It is difficult to predict this spectrum of macrophage phenotypes using our *in vitro* system. However, our study addresses potential mechanisms underlying CAR T cell and PD-L1 blockade alone and in combination. While our studies did not include validation of this combination therapy approach using *in vivo* models, our histological evaluation of tumors in humanized MISTRG mice do confirm increased PD-L1 expression in TAMs following CAR T cell therapy. We previously developed and published an immunocompetent mouse model where we assessed safety and efficacy of PSCA-CAR T cells in murine cancers,[33]. Future studies will evaluate the combination using this syngeneic mouse model. Additionally, future clinical trials to evaluate safety and efficacy of combining CAR T cell therapy and ICB in solid cancers and lymphoma may corroborate our findings.

To our knowledge, this is the first example of a mode of action of ICB by which myeloid cells are directly targeted and depleted specifically in the context of CAR T cell therapy, and this study gives new insights to a mechanism by which PD-L1-negative tumors may benefit from CAR T cell therapy in combination specifically with PD-L1 blockade. The altered phenotypes and depletion of immune-suppressive macrophages in tumors may require both CAR T cells and PD-L1 blockade and warrants further engineering of CAR T cells to secrete PD-L1 blockers and enhance IFNγ signaling to improve anti-tumor responses in TAM-rich solid tumors.

## Materials and Methods

### Cell lines

Human metastatic prostate cancer cell lines DU145 (ATCC HTB-81) and PC-3 (ATCC CRL-1435), human lymphoma cell line Daudi (ATCC CCL-213), and human monocytic leukemia cell line THP-1 (ATCC TIB-202) were cultured in RPMI-1640 (Lonza, 12-115F) containing 10% fetal bovine serum (FBS, Hyclone, SH30070.03) (RPMI+10%FBS). DU145 and PC-3 tumor cells were engineered to express PSCA antigen as previously described,[35]. Human pancreatic cancer cell line HPAC (ATCC CRL-2119) and human breast cancer cell line MDA-MB-231 (ATCC CRM-HTB-26) were cultured in Dulbecco’s Modified Eagle Medium: Nutrient Mixture F-12 (DMEM/F12, Corning, 10-092-CV) containing 10% FBS. MCF-7 (ATCC HTB-22) breast cancer cells were cultured in Dulbecco’s Modified Eagle Medium (DMEM, Gibco, 11960-051) containing 10% FBS, 25 mM HEPES (Irvine Scientific, 9319), and 2mM L-Glutamine (Lonza, 17-605E). Patient-derived metastatic prostate cancer LAPC-9 cells used in vivo were generously provided by the Reiter Lab at UCLA. LAPC9 cells were engineered to express eGFP/firefly luciferase (LAPC-9-eGFP-ffLuc) and maintained as described in previous literature,[35].

### DNA Construction and Lentivirus production

PSCA- and CD19-targeting CARs were designed as previously described, and respective constructs carried truncated CD19 and EGFR as a surrogate marker of transduction,[35, 36]. Lentivirus was manufactured following previously established methods,[35]. In short, lentivirus was generated using 293T cells in T-225 flasks and cultured overnight prior to transfection with packaging plasmids and desired lentiviral backbone plasmid. Supernatants containing lentivirus were collected following 3 to 4 days, filtered, and centrifuged to remove residual cell debris. Lentivirus containing supernatant then underwent incubation with 2mM magnesium and 25U/mL Benzonase endonuclease. Suspended lentivirus was then concentrated by high-speed centrifugation (6080 x g) overnight at 4°C. Lentiviral pellets were resuspended in PBS-lactose solution (4g lactose per 100mL PBS) then aliquoted and stored at -80°C until ready for use. Lentiviral titers were determined using Jurkat cells.

### PBMC and monocyte isolation

Leukapheresis products were obtained from consented research participants (healthy donors) under protocols approved by the City of Hope Internal Review Board (IRB). On the day of leukapheresis, peripheral blood mononuclear cells (PBMC) were isolated by density gradient centrifugation over Ficoll-Paque (GE Healthcare) followed by multiple washes in PBS/EDTA (Miltenyi Biotec).

Monocytes were isolated from freshly collected PBMCs using CD14 antibody-conjugated microbeads and magnetic columns (Miltenyi Biotec) according to the manufacturer’s protocol. CD14^+^ monocytes and CD14^-^ fraction were frozen in CryoStor^®^ CS5 (StemCell Technologies) until processed further.

### T cell lentiviral transduction and *ex vivo* expansion

T cell activation and transduction was performed as described previously,[35]. Briefly, freshly thawed CD14^-^ or whole PBMCs were washed once and cultured in X-VIVO-15 (Lonza) with 10% FBS (complete X-VIVO) containing 100 U/mL recombinant human IL-2 (rhIL-2, Novartis Oncology) and 0.5 ng/mL recombinant human IL-15 (rhIL-15, CellGenix). For CAR lentiviral transduction, T cells were cultured with CD3/CD28 Dynabeads^®^ (Life Technologies), protamine sulfate (APP Pharmaceuticals), cytokine mixture (as stated above), and desired lentivirus at a 0.1-1 multiplicity or infection (MOI) the day following stimulation. Cells were then cultured in and replenished with fresh complete X-VIVO containing cytokines every 2–3 days. After 7 days, beads were magnetically removed, and cells were further expanded in complete X-VIVO containing cytokines to achieve desired cell yield. CAR T cells were positively selected for truncated CD19 using the EasySep™ CD19 Positive Enrichment Kit I or II (StemCell Technologies) (for PSCA-CAR T cells) or positively selected for truncated EGFR using a custom EasySep™ EGFR Positive Enrichment Kit (for CD19-CAR T cells) according to the manufacturer’s protocol. Following further expansion, cells were frozen in CryoStor^®^ CS5 prior to *in vitro* functional assays and *in vivo* therapeutic models. Purity and phenotype of CAR T cells were verified by flow cytometry.

### *In vitro* macrophage differentiation

Primary human M1 and M2 macrophages were differentiated and polarized as previously described,[34]. Briefly, frozen human monocytes were thawed and cultured in cytokine-containing RPMI+10% FBS for 7-10 days. To differentiate M1 macrophages, cells were cultured with GM-CSF (BioLegend, 572903). The media was changed once after 3-5 days to media containing GM-CSF, IFNγ (BioLegend, 570202), LPS (Sigma-Aldrich, L3012-5MG) and IL-6 (BioLegend, 570804). To differentiate M2 macrophages, cells were cultured with M-CSF (BioLegend, Cat: 574804). The media was changed once after 3-5 days to media containing M-CSF, IL-4 (BioLegend, 574004), IL-13 (BioLegend, 571102) and IL-6. All cytokines and LPS were used at 20ng/mL. After differentiation, macrophages were lifted using PBS + 1mM EDTA (PBS-EDTA, Cellgro), and phenotype was assessed by flow cytometry to confirm successful polarization. Cells were counted and used for further studies.

To differentiate and polarize M1 and M2 macrophages from human monocytes THP-1 (ATCC TIB-202), commonly used protocols were adapted,[46, 47]. THP-1 cells were stimulated with phorbol 12-myristate 13-acetate (PMA) for 24 hours and rested for 72 hours in RPMI+10% FBS. Cells were then polarized for 24 hours to M1 or M2 macrophages in the presence of IFNγ and LPS or IL-4 and IL-13, respectively. Cytokines and PMA were used at 20ng/mL. Polarized macrophages were lifted with PBS-EDTA and used for further studies.

### Flow cytometry

For flow cytometric analysis, cells were resuspended in FACS buffer (Hank’s balanced salt solution without Ca^2+^, Mg^2+^, or phenol red (HBSS^−/−^, Life Technologies) containing 2% FBS and 0.5% Sodium Azide. Cells were incubated with primary antibodies for 30 min at 4°C in the dark. Cell viability was determined using 4’,6-diamidino-2-phenylindole (DAPI, Sigma). Flow cytometry was performed on a MACSQuant Analyzer 10 (Miltenyi Biotec), and the data were analyzed with FlowJo software (v10, TreeStar). Antibodies targeting human CD45 (BD Biosciences, 347464), CD137 (BD Biosciences, 555956), CD19 (BD Pharmingen™, 557835), EGFR (BioLegend, 352906), CD80 (BD Biosciences, 340294), CD163 (eBioscience, 17-1639-42), CD206 (BioLegend, 321123), PD-L1 (BD Biosciences, 558065), PD-1 (eBioscience, 47-2799-42), CD33 (BD Biosciences, 340533), HLA-DR (eBioscience, 47-9956-42), and CSF1R (BioLegend 347305) were used for analysis.

### ELISA

IFNγ in supernatant was measured using Human IFNγ ELISA Kit (Invitrogen, 88-7316-88) according to the manufacturer’s protocol. Plates were read at 450 nm using Cytation 5 (BioTek).

### *In vitro* immune-suppression assay

CAR T cells, macrophages, and target tumors were co-cultured in RPMI+10% FBS in the absence of exogenous cytokines in 96-well plates. Cells were plated at an effector:macrophage:target (E:M:T) ratio of 1:5:10 to model prostate cancer with DU145-PSCA cells and lymphoma with Daudi cells. For analysis of the prostate cancer model, supernatant was collected after 3 days for ELISA, and cells were trypsinized and collected for flow cytometry after 6 or 10 days. T cell proliferation was assessed after 10 days, and all other parameters including tumor cell killing, T cell activation and macrophage phenotype were evaluated after 6 days by flow cytometry. The lymphoma model was analyzed after 3 days of culture.

### Generation of CAR T cell-derived conditioned media

PSCA-CAR T cells or untransduced (UTD) controls (5 × 10^3^) were co-cultured with DU145-PSCA cells (5 × 10^4^) for 72 hours, and supernatant was collected and centrifuged at 500 x g for 5 minutes. Cell-free conditioned media (CM) was collected and stored at -80°C. CAR T cell function was validated by flow cytometry, and when it is mentioned, ELISA was performed prior to using the supernatant to determine concentrations of IFNγ.

### Stimulation of macrophages with CAR T cell-derived conditioned media

Differentiated macrophages were plated in RPMI+10% FBS and rested overnight, and CAR or UTD T cell-derived CM collected from tumor cell-T cell co-cultures was applied to stimulate macrophages. Cells were analyzed by flow cytometry after 48 hours. Cell morphology was captured by using BZ-X810 Inverted Microscope (Keyence) or Axio Vert.A1 Inverted Microscope (Zeiss). Atezolizumab (anti-human PD-L1, Tecentriq®, Genentech), Avelumab (anti-human PD-L1, Bavencio®, EMD Serono), Nivolumab (anti-human PD-1, Opdivo®, Bristol Meyers Squibb), and isotype control (bgal-mab12, InvivoGen) were added at the time of stimulation. Anti-IFNγRα (BioLegend, 308610) and isotype control (BioLegend, 400166) were added to culture 2 hours prior to stimulation with CM. Similarly, cells were pre-incubated with small molecule inhibitors for 30 minutes prior to stimulation. Small molecule inhibitors included Fludarabine (STAT1 inhibitor, EnzoALX-480-100-M005), AZD1480 (JAK1 & JAK2 inhibitor, MilliporeSigma, SML1505-5MG), Itacitinib (JAK1 inhibitor, Cayman Chemicals, 27597), Rapamycin (mTOR inhibitor, Cayman Chemicals, 13346), C188-9 (STAT3 inhibitor, Cayman Chemicals, 30928), Akt Inhibitor VIII (AKT inhibitor, MilliporeSigma, 124018-5MG), BAY 11-7082 (NF-κB inhibitor, MilliporeSigma, B5556-10MG), AG490 (JAK2 inhibitor, MilliporeSigma, 658401-5MG), CZC24832 (PI3Kγ inhibitor, MilliporeSigma, SML1214-5MG).

### RT-PCR

RNA was isolated using RNeasy mini kit (Qiagen) or Quick-RNA Microprep Kit (Zymo Research), and RNA concentration was measured using NanoDrop (Thermo Scientific). cDNA was prepared from 0.4-1∼g of total RNA using SuperScript IV reverse transcriptase (ThermoFisher Scientific). Quantitative PCR was performed using SsoAdvanced Universal SYBR Green Supermix (Bio-Rad) on CFX96 Real-Time PCR Detection System (Bio-Rad). The data were analyzed by the comparative threshold method, and gene expression was normalized to *GAPDH*. The following primers were used: *CD274*: forward, GCTGAACGCCCCATACAACA; reverse, TCCAGATGACTTCGGCCTTG and *GAPDH*: forward, TCGGAGTCAACGGATTTGGT; reverse, TTCCCGTTCTCAGCCTTGAC. These primer sets were validated to have a single melting curve and amplification efficiency of 2.

### RNA sequencing

Macrophages were stimulated with CAR or UTD T cell-derived CM collected from tumor cell-T cell co-cultures. After 8 hours, cells were lysed using RNA Lysis Buffer (Zymo Research, R1060-1-50), and RNA was isolated according to the manufacturer’s protocol. Libraries for stranded poly(A) RNA-seq were created using the KAPA mRNA HyperPrep kit (Roche).

Sequencing of 51 bp single-end reads was performed using a HiSeq2500 regular run. Base calling (de-multiplexing samples between and within labs by 6 bp barcodes, from a 7 bp index read) was performed using bcl2fastq v2.18. Reads were aligned against the human genome using TopHat2,[48]. Read counts were tabulated using htseq-count,,[49] with UCSC known gene annotations,[50]. Change values were calculated from fragments per kilobase per million (FPKM) reads normalized expression values, which were also used for visualization (following a log2 transformation),[51]. Aligned reads were counted using GenomicRanges,[52]. GSEA was run on log2(FPKM + 0.1) expression values, with upregulated enrichment results for GO Biological Process categories in MSigDB,[53-55].

### Animal Experiments

All animal experiments were performed under protocols approved by the City of Hope Animal Care and Use Committee (IACUC). MISTRG mice were obtained through MTA from Regeneron Pharmaceuticals and housed and bred at City of Hope. 3-6 week old MISTRG mice were sublethally irradiated (100cGy, J.L. Shepherd Mark I Cs-137 irradiator) 6-12 hours prior to engraftment of human adult G-CSF mobilized CD34^+^ cells (2.5 × 10^5^) via intravenous injection. Human adult G-CSF mobilized CD34^+^ cells and autologous PBMCs were purchased from HemaCare, and autologous PBMCs were used to manufacture CAR and UTD T cells used for adoptive cell transfer (ACT). DU145-PSCA cells (2.5-5 × 10^5^) were engrafted subcutaneously (s.c.), and tumor growth was monitored by biweekly caliper measurement. For an orthotopic intratibial model, LAPC-9-eGFP-ffLuc cells (1.5 × 10^5^) were engrafted into the intratibial space (i.ti.), and tumor growth was monitored by biweekly non-invasive bioluminescence imaging (Lago-X, Accela). For non-invasive flux imaging, mice were injected intraperitonially with 150 mL D-luciferin potassium salt (Perkin Elmer) suspended in PBS at 4.29 mg/mouse. Flux signals were analyzed with Aura imaging software (Spectral Instruments Imaging). Mice received ACT of CAR or UTD T cells (1 × 10^6^) when DU145-PSCA s.c. reach ~150mm^3^ or 14 days after LAPC-9-eGFP-ffLuc i.ti. engraftment. Tumors were harvested 7 days following ACT for histology.

### Immunohistochemistry and Immunofluorescent Staining

Collected mouse tissue was fixed in 4% paraformaldehyde (4% PFA, Boston BioProducts) and stored in 70% ethanol until processed further. Tissue embedding, sectioning, H&E and IHC staining were performed by the Research Pathology Core at City of Hope.

Immunofluorescent staining of tissue was completed on paraffin embedded tissue. In brief, paraffin sections were deparaffinized and rehydrated, and antigens were retrieved in citrate-based antigen unmasking solution (Vector Lab, H-3300-250) for 10 minutes at 120°C using an autoclave. Samples were rehydrated, permeabilized with 0.1% Triton X-100 for 30 minutes at room temperature and blocked with 5% normal donkey serum (NDS) for 45 minutes prior to immunostaining. Tissue was incubated with rabbit anti-human CD68 (1:200, Cell Signaling Technology, 76437T) and goat anti-human PD-L1 (1:50, Leinco Technologies, B560) at 4°C overnight and washed in PBS+0.1% Tween 20 for 5 min three times. Tissue was incubated with secondary antibodies donkey anti-rabbit IgG, AlexaFluor 488 (1:1000, Invitrogen, A-21206) and donkey anti-goat IgG, AlexaFluor 546 (1:1000, Invitrogen, A-11056) for 1 hour at room temperature, washed in PBS+0.1% Tween 20 for 5 min three times and mounted with mounting media containing DAPI (Vector Laboratories). Fluorescent images were captured using BZ-X810 Inverted Microscope (Keyence).

Double IHC was performed by the Research Pathology Core at City of Hope. Staining was performed on Ventana Discovery Ultra (Ventana Medical Systems, Roche Diagnostics, Indianapolis, USA) IHC Auto Stainer, and mouse anti-human CD68 (Dako, M087601-2) and rabbit anti-human PD-L1 (Ventana, 790-4905) were used at 1:100. Briefly, the slides were loaded on the machine, deparaffinization, rehydration, endogenous peroxidase activity inhibition and antigen retrieval were first performed. Two antigens were sequentially detected and heat inactivation was used between the two antigen detection steps to prevent any potential cross-reactivities. Following the first primary antibody (PD-L1) incubation, DISCOVERY anti-Rabbit NP and DISCOVERY anti-NP-AP were incubated, and stains were visualized with DISCOVERY Yellow Kit. Following the heat inactivation, the second primary antibody (CD68) was incubated, DISCOVERY anti-Rabbit HQ and DISCOVERY anti-HQ-HRP were added, and stains were visualized by DISCOVERY Teal Kit. The slides were then counterstained with hematoxylin (Ventana) and coverslipped. Slides were scanned by using NanoZoomer 2.0HT (Hamamatsu).

### Statistical Analysis

Data are presented as mean ± standard error of the mean (SEM) unless otherwise stated. Statistical comparisons between groups were performed using the unpaired two-tailed Student’s *t* test to calculate *p* values, unless otherwise stated.

## Supplemental Information

Supplemental figures and legends are available and included as separate document.

## Acknowledgments

We thank the staff members of the following cores at the Beckman Research Institute at City of Hope Comprehensive Cancer Center: Animal Facility, Pathology, Small Animal Imaging, and Light Microscopy for their excellent technical assistance. We thank Charles War-den and Dr. Xiwei Wu of the Integrative Genomics Core for their technical assistance in RNAseq analysis. We would also like to thank Catalina Martinez at City of Hope for contributions to manuscript editing.

## Funding

Research reported in this publication was supported by a Prostate Cancer Foundation Young Investigator Award (PI: Y.Y.), a T.J. Martell Foundation Cancer Research Grant (PI: S.J.P.), and a Department of Defense Idea Development Award (S.J.P., W81XWH-17-1-0208). Work performed in the Pathology Core and Small Animal Imaging Core was supported by the National Cancer Institute of the National Institutes of Health under grant number P30CA033572. The content is solely the responsibility of the authors and does not necessarily represent the official views of the National Institutes of Health.

## Author contributions

S.J.P. and Y.Y. provided conception and construction of the study. S.J.P, S.J.F., T.B.D., X.W., J.C.Z., V.D.J., N.L., R.N., K.O., J.G., and Y.Y. provided design of experimental procedures, data analysis, and interpretation. Y.Y., J.G., and K.O., performed experiments. Y.Y. and S.J.P. wrote the manuscript. C.M. and J.G. assisted in writing the manuscript. S.J.P. supervised the study. All authors reviewed the manuscript.

## Competing interests

S.J.P. and S.J.F. are scientific advisors to and receive royalties from Mustang Bio. S.J.P. is a scientific advisor and/or receives royalties from Imugene Ltd., Bayer, and Adicet Bio. All other authors declare that they have no competing interests.

