## Supplementary figures and images for "PD-L1 blockade restores CAR T cell activity through IFNγ-regulation of CD163+ macrophages"

### Supplemental Figures

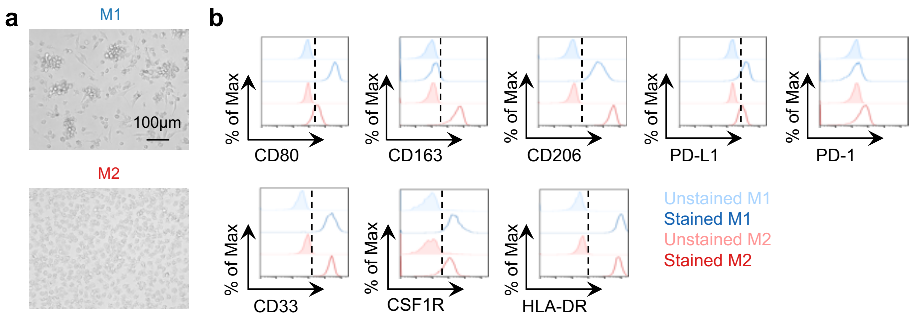

Figure S2

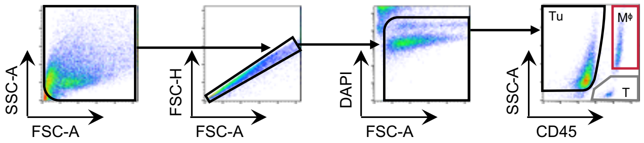

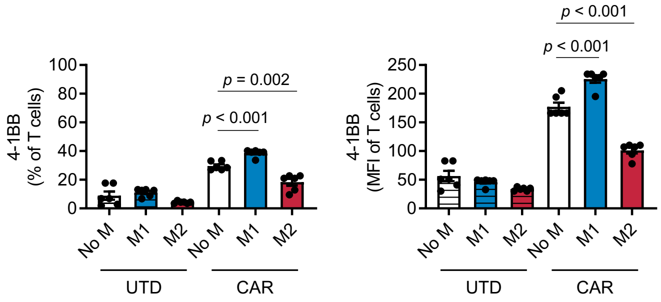

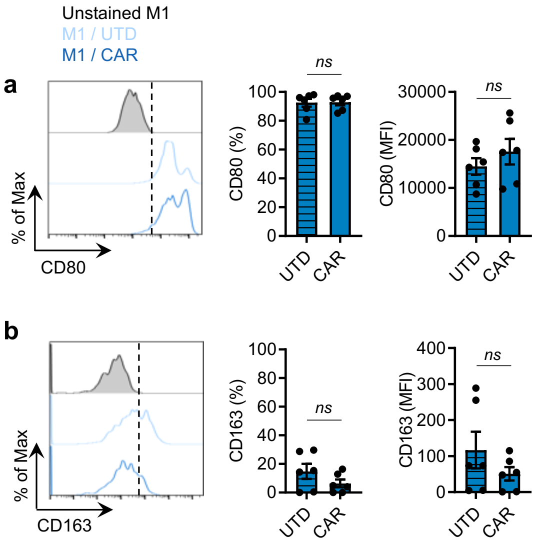

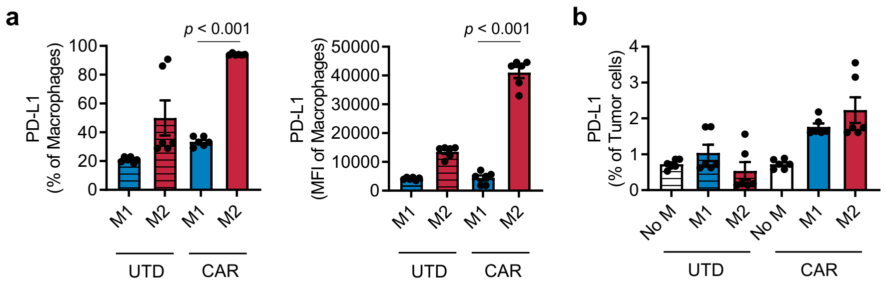

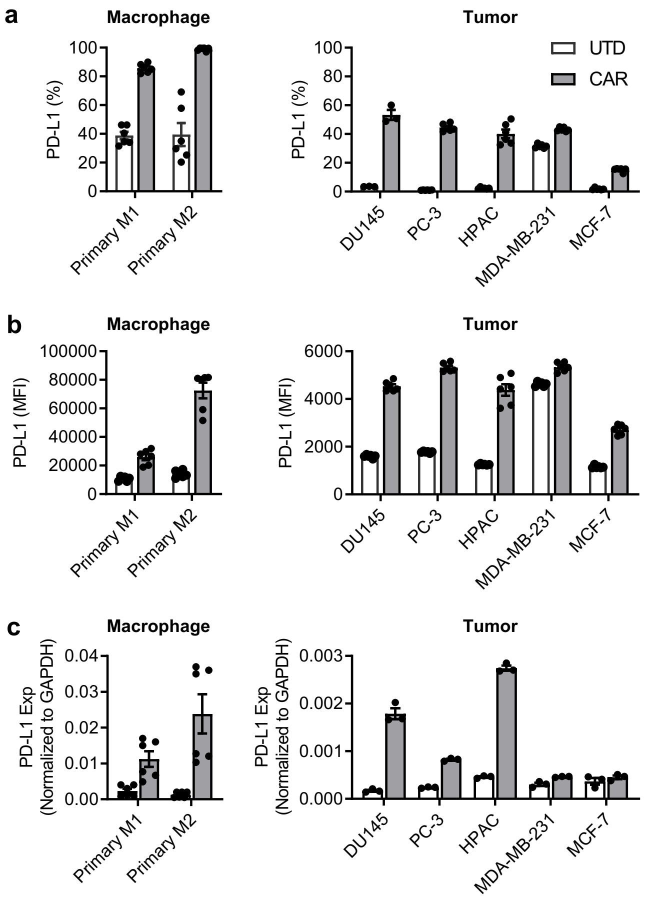

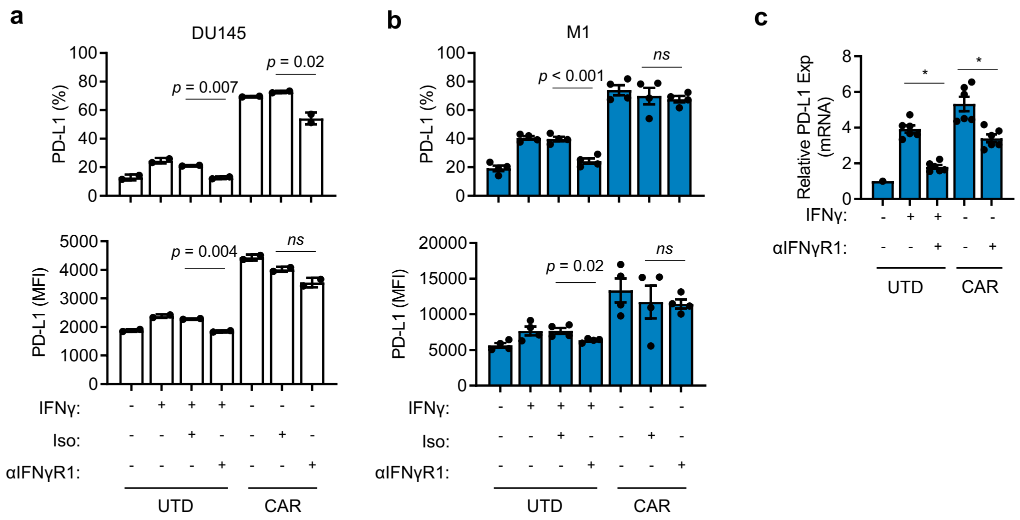

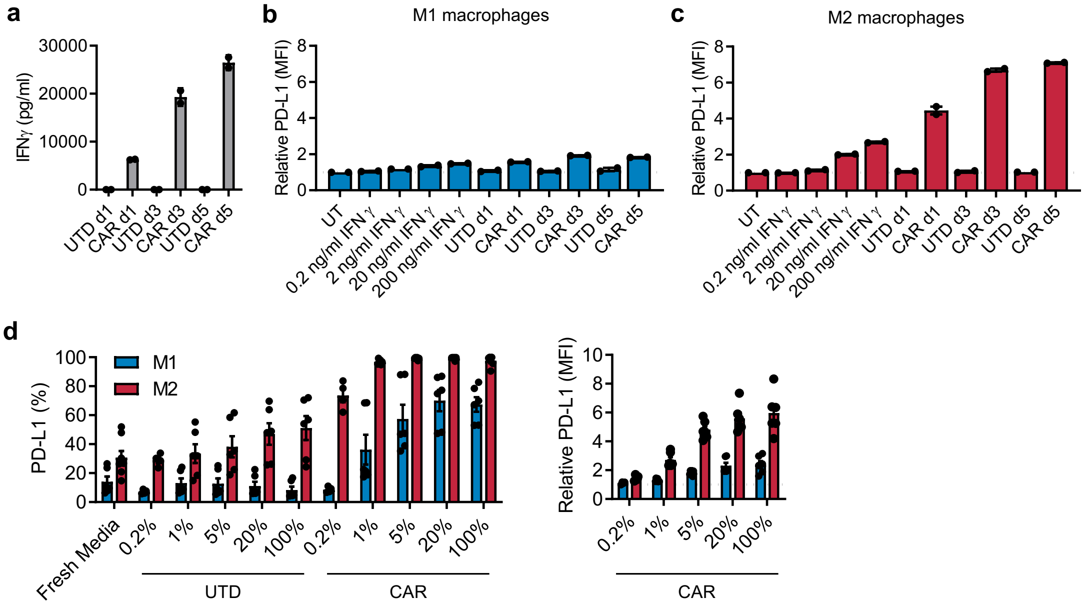

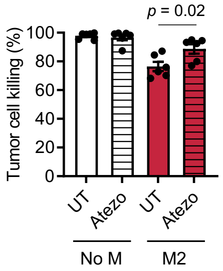

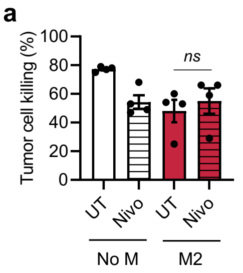

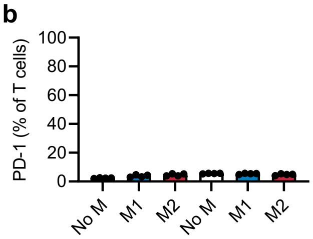
