## Supplemental Figure Legends for "PD-L1 blockade restores CAR T cell activity through IFNγ-regulation of CD163+ macrophages"

Yamaguchi et al.

**Figure S1: *In vitro* differentiation and polarization of human M1 and M2 macrophages.** (a) Images of polarized M1 and M2 macrophages captured by microscopy. (b) Cell surface marker expression assessed by flow cytometry. CD80 is a M1 macrophage marker, and CD163 and CD206 are M2 macrophage markers.

**Figure S2: Gating strategies used for flow cytometry analysis of the immune-suppression assay.** DU145-PSCA tumor cells, T cells and macrophages were separated by representative gating strategies.

**Figure S3: M2 macrophages suppress activation of CD19-CAR T cells targeting CD19+ Daudi lymphoma cells.** 4-1BB T cell activation was evaluated by flow cytometry in the immune-suppression assay modeling lymphoma TME.

**Figure S4: CAR T cells do not alter M1 macrophage phenotypes.** Cell surface expression of CD80 (a) and CD163 (b) in M1 macrophages evaluated by flow cytometry following 48 hour stimulation with CAR T cell-derived CM.

**Figure S5: PD-L1 induction in M2 macrophages in the immune-suppression assay of CD19-CAR T cells targeting CD19+ Daudi lymphoma cells.** (a) Frequency and abundance of PD-L1 induced in M1 or M2 macrophages by CAR T cells. (b) Lack of PD-L1 expression or induction in Daudi tumor cells.

**Figure S6: PD-L1 in macrophages and various tumors induced by CAR T cell-derived CM.** (a, b) Frequency (a) and abundance (b) of PD-L1 expression assessed by flow cytometry 48 hours after stimulation with CAR T cell-derived CM. (c) Induction of PD-L1 mRNA evaluated by RT-PCR following 6 hour stimulation with CAR T cell-derived CM.

**Figure S7: Inhibition of IFN**γ **signaling does not block PD-L1 induction.** Anti-IFNγR1 antibody was used to block IFNγ signaling in the presence or absence of recombinant IFNγ or CAR T cell-derived CM. (a, b) Frequency and abundance of PD-L1 protein in DU145 tumor cells (a) and M1 macrophages (b) were evaluated by flow cytometry 48 hours after stimulation with CAR T cell-derived CM. (c) Induction of PD-L1 mRNA in M1 macrophages was quantified by RT-PCR following 6 hour stimulation with CAR T cell-derived CM.

**Figure S8: CAR T cell-derived cytokines are a potent inducer of PD-L1 expression.** (a) Concentration of IFNγ in CAR T cell-derived CM collected from tumor killing assay. CM was collected after 1, 3, or 5 days in the assay, and IFNγ was measured by ELISA. (b, c) Induction of PD-L1 expression with varying concentration of recombinant IFNγ in M1 (b) and M2 (c) macrophages. Induction in the abundance of PD-L1 is shown relative to an untreated condition of respective cell types. CAR T cell-derived CM with known concentration of IFNγ (a) was used in this assay as reference to recombinant IFNγ. (d) PD-L1 induction of M1 and M2 macrophages by serial dilutions of CAR T cell-derived CM. Frequency and abundance of PD-L1 protein were assessed by flow cytometry. Relative abundance was calculated by normalizing to the abundance of PD-L1 in respective dilutions of UTD T cell-derived CM.

**Figure S9: PD-L1 blockade restores CD19-CAR T cell function inhibited by suppressive M2 macrophages in the Daudi lymphoma model.** CAR T cell-mediated killing of Daudi tumor cells in the presence or absence of Atezo.

**Figure S10: Blocking PD-1 in the immune-suppression assay does not restore CAR T cell function.** (a) PD-1 expression in PSCA-CAR T cells in the immune-suppression assay. (b) Tumor killing by CAR T cells evaluated in the presence of anti-PD-1 antibody, Nivolumab (Nivo).
